# The long-term effects of genomic selection: Response to selection, additive genetic variance and genetic architecture

**DOI:** 10.1101/2021.03.16.435664

**Authors:** Yvonne C.J. Wientjes, Piter Bijma, Mario P.L. Calus, Bas J. Zwaan, Zulma G. Vitezica, Joost van den Heuvel

**Author notes:** Author information: Yvonne Wientjes, Wageningen University & Research, Animal Breeding and Genomics, P.O. box 338, 6700 AH Wageningen, the Netherlands.

## Abstract

Genomic selection has revolutionized genetic improvement in animals and plants, but little is known of its long term effects. Here we investigate the long-term effects of genomic selection on the change in the genetic architecture of traits over generations. We defined the genetic architecture as the subset, allele frequencies and statistical additive effects of causal loci. We simulated a livestock population under 50 generations of phenotypic, pedigree, or genomic selection for a single trait, controlled by either only additive, additive and dominance, or additive, dominance and epistatic effects. The simulated epistasis was based on yeast data. The observed change in genetic architecture over generations was similar for genomic and pedigree selection, and slightly smaller for phenotypic selection. Short-term response was highest with genomic selection, while long-term response was highest with phenotypic selection, especially when non-additive effects were present. This was mainly because the loss in genetic variance and in segregating loci was much greater with genomic selection. Compared to pedigree selection, genomic selection lost a similar amount of the genetic variance but maintained more segregating loci, which on average had lower minor allele frequencies. For all selection methods, the presence of epistasis limited the changes in allele frequency and the fixation of causal loci, and substantially changed the statistical additive effects over generations. Our results show that non-additive effects can have a substantial impact on the change in genetic architecture. Therefore, non-additive effects can substantially impact the accuracy and future genetic gain of genomic selection.

## INTRODUCTION

Animal breeding has substantially increased the performance of livestock populations over the last century (Hill and Kirkpatrick 2010; Hill 2016). This has been achieved by selecting the genetically best performing individuals to produce the next generation based on own performance and/or performances of relatives. Despite the strong selection, these pedigree-based selection methods have proven to be sustainable; genetic variation and rates of genetic gain have been stable for many generations in several animal and plant species, both in commercial breeding programs and experimental selection lines (Beniwal *et al*. 1992; Dudley and Lambert 2003; Havenstein *et al*. 2003a, b).

Recently, genomic selection has revolutionized animal breeding (Meuwissen *et al*. 2001; Meuwissen *et al*. 2016). Within genomic selection, several thousands of DNA markers covering the whole genome are used to identify the genetically best animals. In some breeding programs, genomic selection has doubled the annual rate of genetic gain compared to classical pedigree selection (Schaeffer 2006; García-Ruiz *et al*. 2016). Arguably, genomic selection enables selection for low-heritability traits (Calus *et al*. 2008; Wolc *et al*. 2011) and traits that are difficult or expensive to measure (Goddard and Hayes 2009; Daetwyler *et al*. 2012; Calus *et al*. 2013), for which classical selection is generally difficult. These properties have resulted in a rapid uptake of genomic selection in animal breeding programs worldwide (Hayes *et al*. 2009; Knol *et al*. 2016; Meuwissen *et al*. 2016; Wolc *et al*. 2016).

The accuracy and, thereby, genetic gain of genomic selection are affected by the genetic architecture of the traits (Daetwyler *et al*. 2010; Hayes *et al*. 2010; Wientjes *et al*. 2015), that is the set of causal loci underlying the trait, their frequencies, and their statistical additive effects. The genetic architecture is largely unknown for most traits, including those under selection in breeding programmes, but is known to evolve over time as a result of new mutations and changing allele frequencies due to selection and drift (Wright 1931; Robertson 1960; Falconer and Mackay 1996; Hansen *et al*. 2006; Le Rouzic and Carlborg 2008; Hill and Kirkpatrick 2010; Hill 2016). When interactions are present within (dominance) or between (epistasis) loci, the statistical additive effects (also known as allele substitution effects) depend on the allele frequency at the locus itself as well as of those at interacting loci. This means that part of the functional dominance and epistatic effects contribute to additive genetic variation, with their total contribution depending on the allele frequencies (Barton and Turelli 2004; Hill *et al*. 2008; Mäki-Tanila and Hill 2014). Although interactions between loci are known and common (Carlborg and Haley 2004; Carlborg *et al*. 2006; Flint and Mackay 2009; Huang *et al*. 2012), not much is known about their interaction network or how those interactions convert into genetic variance components or change over generations as a result of drift or selection. The genetic interaction network is so far most intensively studied in yeast, where 90% of the loci associated with a trait were found to be involved in at least one interaction, with only few interactions for most of the loci, and a lot of interactions for only a few loci (Tong *et al*. 2004; Boone *et al*. 2007; Costanzo *et al*. 2016). Boone *et al*. (2007) and Mackay (2014) argue that it is likely that this genetic interaction network is similar in other species such as livestock and human as well.

We hypothesize that genomic selection accelerates the change in genetic architecture of traits across generations, which can affect long-term genetic gain. The reason is not only that genomic selection is more effective, but also related to the distribution of the selection pressure across the genome. Classical selection methods based on pedigree relationships implicitly weigh effects of alleles independently of allele frequency or effect size and distribute selection pressure evenly across the genome (Goddard 2009). This is in contrast to genomic selection methods that put less weight on rare alleles (Goddard 2009; Bijma 2012). Genomic selection methods, therefore, more strongly select on genomic regions surrounding loci with a large contribution to the additive genetic variance and may significantly increase the change in allele frequency at those loci (Heidaritabar *et al*. 2014). Therefore, genomic selection may substantially accelerate the rate of genetic gain in the short-term, but by ignoring regions with a smaller contribution to additive genetic variance, genomic selection increases the risk of losing rare favourable alleles or may fail to increase frequency of such alleles (Jannink 2010; Liu *et al*. 2015; De Beukelaer *et al*. 2017). The loss of rare favourable alleles reduces genetic variation and genetic gain in the long term (Goddard 2009), and limits the potential to accommodate changes in desirable phenotypes in the future. However, currently, these expectations have not been investigated in detail or tested in breeding populations.

Therefore, the aim of this study is to investigate the long-term effects of genomic selection on the genetic architecture of traits. Using simulations, genomic selection will be compared to phenotypic and pedigree selection. We will investigate the impact of those selection methods on the rate of genetic gain, the loss in genetic variance and the change in genetic architecture for 50 generations of selection. Those results will give us more insight on the long-term evolution of the genetic architecture and genetic variation of traits under different selection methods.

## MATERIALS AND METHODS

### Simulated population

We simulated a livestock population under 50 generations of selection. As a first step, we constructed a historical population in which selection was absent and mating was at random, using the QMSim software (Sargolzaei and Schenkel 2009). The first 2000 generations (generation -3050 to -1050) consisted of 1500 individuals, after which the size of the population gradually decreased to 100 over 500 generations (generation -1050 to -550) to resemble a bottleneck in the population and to generate linkage disequilibrium. This was followed by a gradual increase in population size to 1500 over 500 generations (generation -550 to -50). From the last historical generation (generation -50), 100 females and 100 males were randomly sampled, and their genotypes were the input for our own developed Fortran program. Those individuals were randomly mated (mating ratio 1:1) with a litter size of 10 (5 females and 5 males). In each of the next 50 discrete generations, 100 females and 100 males were randomly sampled and mated to build up mutation-drift equilibrium (generation -50 to 0), using a selected proportion of 0.2. Generation 0 formed the base population for the 50 generations of selection. In the following generations, we used truncation selection to select the best 100 females and 100 males that were randomly mated using a mating ratio of 1:1 and a litter size of 10 (5 females and 5 males), resulting in a selected proportion of 0.2 for both females and males. Five selection methods were used which will be explained later.

### Genome

The simulated genome contained 10 chromosomes of 100 cM each. The number of recombination events per chromosome was sampled from a Poisson distribution with on average one recombination per chromosome and a random allocation of the recombination location on the chromosome.

In the historical population, 200,000 randomly spaced bi-allelic loci per chromosome were simulated with a recurrent mutation rate of 5 * 10^−5^. The population structure and mutation rate resulted in a U-shaped allele frequency distribution of the loci in the historical population. In the last historical generation, 2000 segregating loci were randomly selected to become causal loci. Another set of 20,000 segregating loci were selected as marker, by selecting 200 loci from each of 100 equally sized bins based on allele frequency. This resulted in a uniform allele frequency distribution of the markers, reflecting the ascertainment bias on commercial marker chips (Matukumalli *et al*. 2009; Ramos *et al*. 2009; Groenen *et al*. 2011).

After the historical population, the number of mutations per individual was sampled from a Poisson distribution with an average of 0.6. This procedure resulted in a mutational variance of 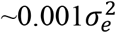 under our simulated additive model (as explained later), as is often observed in real populations (Hill 1982; Houle *et al*. 1996; Lynch and Walsh 1998). As loci for the mutations, we used 4000 loci that did not segregate in the last generation of the historical population and were randomly sampled from all possible loci. The loci and effects of the mutations were recycled to limit the computational requirement. In each generation, a locus was drawn from the potential loci that did not segregate at that moment, while maximizing the time between two mutations at the same locus. As such, each of the 4000 loci was used on average once in every 6-7 generations. We believe that recycling the same mutations does not impact the results of our study, because the vast majority of the mutations are lost in the first generation due to drift, which is unrelated to their effect.

### Genotypic and phenotypic values

Three genetic models were used to simulate phenotypic values; a model with only additive effects (A), a model with additive and dominance effects (AD), and a model with additive, dominance and epistatic effects (ADE). Functional (or biological) additive and dominance effects were assigned to all 2000 causal loci and the 4000 loci for mutations in the last historical generation. At the same time, epistatic effects were assigned to 90% of those loci, as was observed to be the case in yeast data (Costanzo *et al*. 2016).

Functional additive effects (a) were sampled from a normal distribution with mean 0 and standard deviation 1. Functional dominance effects (d) were simulated proportional to the additive effect; we first sampled for each locus a dominance degree (*dd*) from a normal distribution with mean 0.2 and standard deviation 0.3, and subsequently computed the dominance effect of locus *i* as *di=ddi|ai|*. This resulted in mostly positive dominance effects, with a bit of overdominance, as was empirically observed in pigs (Bennewitz and Meuwissen 2010).

Only pairwise epistatic effects were simulated, because higher-order interactions have little effect on the phenotypic values when the allele frequency distribution is U-shaped. The number of interactions per locus was sampled using the interaction network found between the ∼6000 genes in yeast (Stark 2006; Costanzo *et al*. 2016), with many loci with few interactions and few loci with many interactions (Figure 1). This was done by creating an interaction matrix from the network in yeast, with elements of 1 when loci interacted and 0 otherwise. From this matrix, columns and corresponding rows were selected for all loci with an interaction. For the interaction between locus *i* and *j*, nine epistatic degrees (*ε*) were independently sampled from a normal distribution with mean 0 and standard deviation 0.45, one for each of the nine possible two-locus genotype combinations. Those *ε* were used to create nine epistatic effects (*e*) for each interaction as 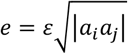 (Table 1), resulting in larger epistatic effects for loci with a larger additive effect. This way of simulating epistatic effects resulted in all types of epistasis, i.e., additive by additive, additive by dominance, and dominance by dominance. However, as a result of simulating the epistatic effects in this random way, part of the simulated epistatic effect represent a functional additive or dominance effect (Table 1). For computing functional additive, dominance and epistatic variance components, we first redistributed the simulated epistatic effects in the correct underlying functional effects.

**TABLE 1.**
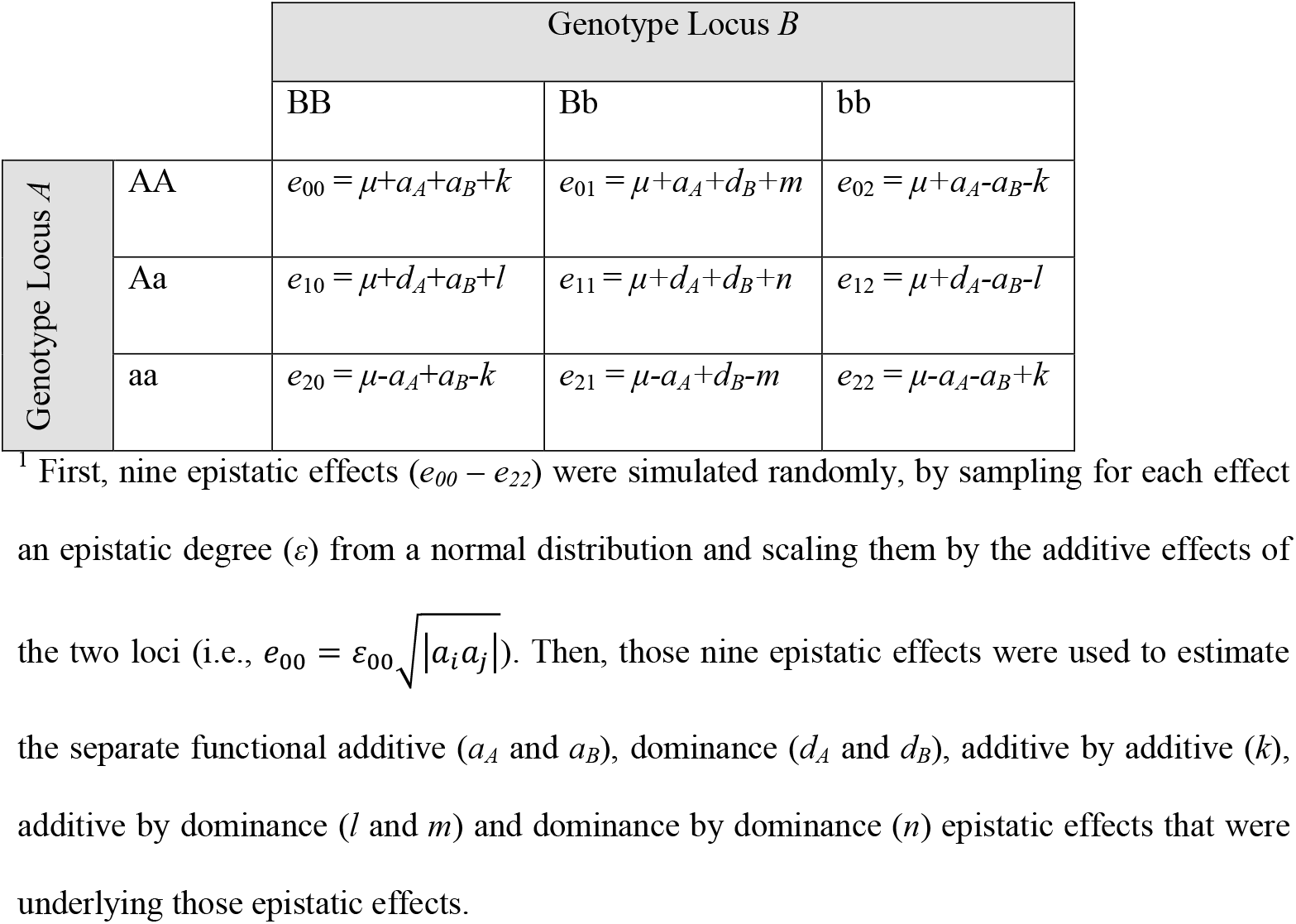
Simulated epistatic model for two-locus interactions^1^

|  |  | Genotype Locus *B* |  |  |
| --- | --- | --- | --- | --- |
|  |  | BB | Bb | bb |
| Genotype Locus *A* | AA | $e_{00} = \mu + a_A + a_B + k$ | $e_{01} = \mu + a_A + d_B + m$ | $e_{02} = \mu + a_A - a_B - k$ |
| | Aa | $e_{10} = \mu + d_A + a_B + l$ | $e_{11} = \mu + d_A + d_B + n$ | $e_{12} = \mu + d_A - a_B - l$ |
| | aa | $e_{20} = \mu - a_A + a_B - k$ | $e_{21} = \mu - a_A + d_B - m$ | $e_{22} = \mu - a_A - a_B + k$ |

<sup>1</sup> First, nine epistatic effects ( $e_{00} - e_{22}$ ) were simulated randomly, by sampling for each effect an epistatic degree ( $\varepsilon$ ) from a normal distribution and scaling them by the additive effects of the two loci (i.e., $e_{00} = \varepsilon_{00} \sqrt{|a_i a_j|}$ ). Then, those nine epistatic effects were used to estimate the separate functional additive ( $a_A$ and $a_B$ ), dominance ( $d_A$ and $d_B$ ), additive by additive ( $k$ ), additive by dominance ( $l$ and $m$ ) and dominance by dominance ( $n$ ) epistatic effects that were underlying those epistatic effects.

**FIGURE 1.**
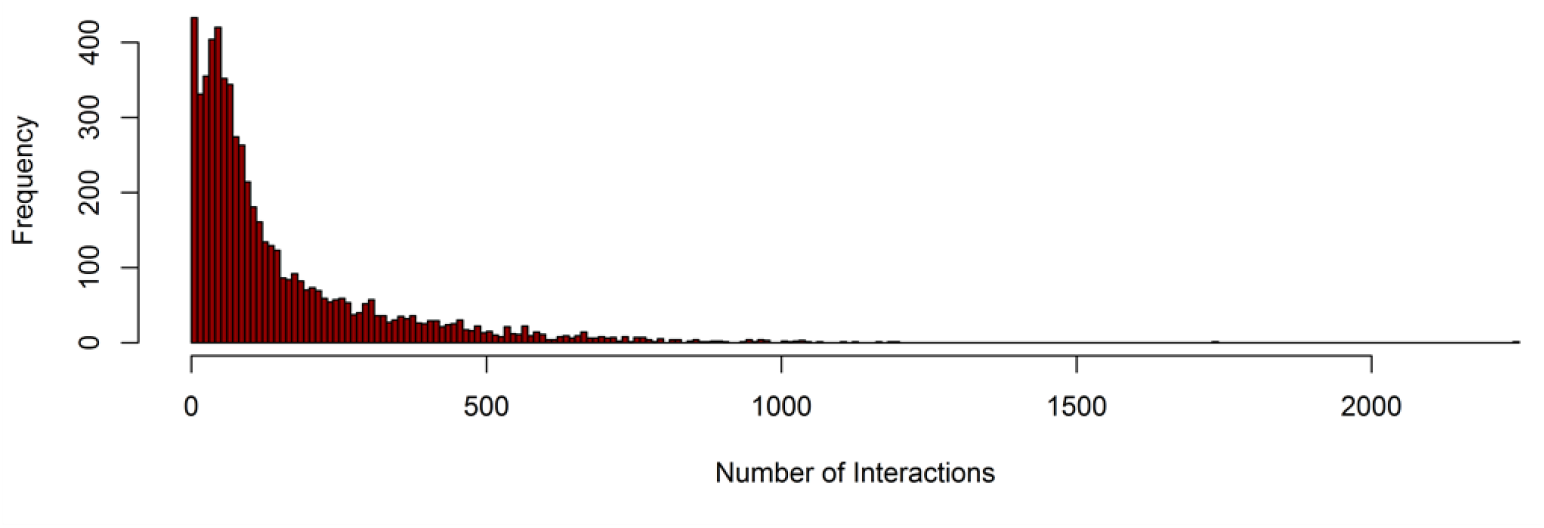
Histogram of the number of interactions per causal locus.

The functional genetic effects were combined with the genotypes of the individuals to calculate total genetic values. For each individual, a residual term was sampled from a normal distribution with mean zero and standard deviation equal to the square root of 1.5 times the variance in total genetic values in the generation in which functional effects were assigned, resulting in a broad sense heritability of 0.4.

### Statistical effects

The natural and orthogonal interaction approach (NOIA) (Álvarez-Castro and Carlborg 2007; Vitezica *et al*. 2017) was applied in each generation to compute statistical additive and dominance effects based on the functional additive, dominance, and epistatic effects of all causal loci (the 2000 segregating causal loci and the 4000 loci for mutations) and their allele frequencies (Duenk *et al*. 2020). For each locus *i*, the part of the dominance effect that is statistically additive was calculated as *(1-2pi)di*, where *p*_*i*_ is the frequency of the focal allele (i.e., allele A for locus A in Table 1). For each interaction between loci *i* (with alleles a and A) and *j* (with alleles b and B) the functional epistasis is converted into statistical additive and statistical dominance effects. These statistical effects were computed from three components: 1) a vector **y** with functional epistatic effects, **y′** = [*e*_00_ *e*_10_ *e*_20_ *e*_01_ *e*_11_ *e*_21_ *e*_02_ *e*_12_ *e*_22_], 2) a 9×9 diagonal matrix **D** with the expected frequencies of the two-locus haplotypes, assuming that loci segregate independently, and 3) a 9×9 matrix **W** with the mean and orthogonal contrasts for the two loci, constructed as **W** *=* **W**_*i*_ ⨂ **W**_*j*_ with

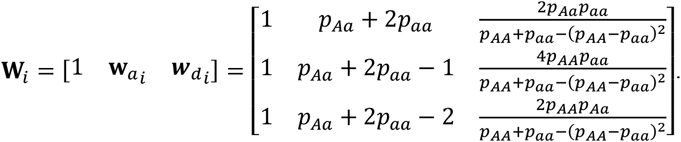

The statistical effects followed from

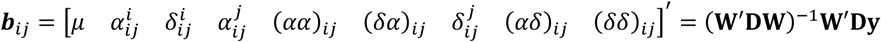

Note that the NOIA model was run separately for each set of interacting loci thereby only considering the functional interaction effects and not the functional additive and dominance effects. Therefore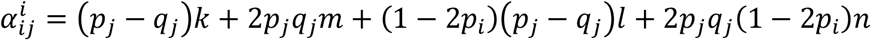, and 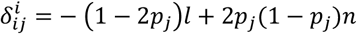; where *k, l, m* and *n* are the additive by additive, dominancy by additive, additive by dominance and dominance by dominance functional epistatic effects (Table 1).

The total statistical additive effect of locus *i* was calculated as

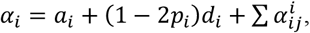

and the total statistical dominance effect as

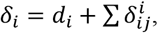

where the summation was taken across all interactions that involved locus *i*.

The statistical additive effect was used to compute the total additive genetic value (i.e., true breeding value) across all loci *<u>i</u>* of each individual as 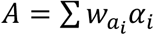, with

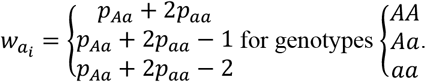

In the same way, the statistical dominance effect was used to compute the total dominance deviation across all loci *i* of each individual as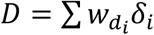, with

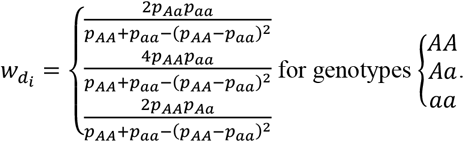

By definition, the variance in *A* across all individuals is the additive genetic variance, and the variance in *D* across all individuals is the dominance genetic variance. The total genetic variance minus the additive and dominance variance is the epistatic variance.

### Selection methods

Five methods were used to select the sires and dams of the next generation. As a base line for comparison, the first method randomly selected the parents (RANDOM) and was meant to capture the impact of drift alone. The second method selected the individuals with the highest phenotypic values to become the parents of the next generation (MASS). The third method selected individuals with the highest estimated breeding values using a pedigree Best Linear Unbiased Prediction (BLUP) model that included own performance information of the selection candidates (PBLUP_OP). The fourth and fifth method selected individuals with the highest genomic estimated breeding values from a genomic BLUP model that either included own performance information of the selection candidates (GBLUP_OP) or not (GBLUP_NoOP).

Breeding value estimation for the last three methods was performed using the MTG2 software (Lee and van der Werf 2016). Breeding values were estimated simultaneously with estimating the variance components, using the phenotypic information of the previous three generations and for PBLUP_OP and GBLUP_OP also phenotypic information of the generation itself. The PBLUP method used a relationship matrix based on a pedigree that included all individuals from the generation itself and the previous eight generations. The GBLUP methods used a relationship matrix based on marker genotypes of the generation itself and the previous three generations, computed using method 1 of VanRaden (VanRaden 2008) with allele frequencies estimated from the genotype data of those generations. The model for breeding value estimation included a fixed mean, a random additive genetic effect, a random litter effect, and a residual. The random litter effect was included to capture the resemblance between full sibs due to non-additive genetic effects, which could otherwise create bias in the estimated breeding values. Even though dominance and epistatic effects were simulated, these were not included in the breeding value estimation model, because additive models are generally used in breeding programs and only the breeding value is transmitted to the offspring.

### Comparing genetic models and selection methods

The three genetic models (A, AD and ADE) and five selection methods were compared based on their accuracy of selection, phenotypic trend, additive genetic variance, additive genic variance (calculated as the sum of 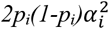 across all causal loci *i*), heterozygosity, average minor allele frequency (MAF), and number of segregating causal loci over the 50 generations of selection. The accuracy of selection was calculated as the correlation between the true breeding values and estimated breeding values. For each of the 15 scenarios, 20 replicates were simulated.

One of our main aims was to evaluate how fast the genetic architecture of the trait changed due to selection. The genetic architecture can change because 1) The subset of loci affecting the trait changes due to new mutations and loci becoming fixed, 2) The allele frequencies of those loci change, which can result in changes in the proportion of the additive genetic variance explained by each locus, or 3) The statistical additive effects of the loci change as a result of allele frequency changes and non-additive effects, which can also change the proportion of additive genetic variance explained by a locus.

We defined three criteria that each reflected one of those mechanisms, namely: 1) The Jaccard index for the segregating causal loci, 2) The correlation in allele frequencies at those loci between generations, and 3) The correlation in statistical additive effects at those loci between generations. For the first criterium, we calculated the Jaccard index (Jaccard 1908) between generation 0 (before selection) and each of the generations after selection as the number of overlapping segregating loci divided by the total number of segregating loci in the two generations. For the second criterium, we calculated the correlation between allele frequencies of generation 0 and each of the generations after selection, using only the loci that segregated in generation 0 and remained segregating. For the third criterium, we calculated the correlation between statistical additive effects of generation 0 with a generation after selection, again including only loci that remained segregating from generation 0 onwards.

## Data availability

Supplemental file 1, contains the QMSim input file, Fortran programs and seeds used to select the markers and causal loci, to simulate functional effects and genotypes and phenotypic values of new generations, and the interaction matrix used to simulate epistatic effects.

## RESULTS

### Properties of simulated population

The allele frequency distribution of the segregating causal loci was strongly U-shaped (Supplemental file 2, Figure S2.1) and comparable to the allele frequency distribution in sequence data of livestock populations (Daetwyler *et al*. 2014; Eynard *et al*. 2015; Heidaritabar *et al*. 2016; Bolormaa *et al*. 2019). In the RANDOM scenario, where no selection was performed, the allele frequency pattern remained similar over generations, indicating that the population was approximately in mutation-drift equilibrium. Moreover, the linkage disequilibrium pattern in the population (Supplemental file 2, Figure S2.2) was similar to that found in livestock populations (Andreescu *et al*. 2007; Badke *et al*. 2012; Veroneze *et al*. 2013). This indicates that the effective population size of the simulated population was comparable to that in real livestock populations that are in the range of 40-130 (Welsh *et al*. 2010; Uimari and Tapio 2011; Wientjes *et al*. 2013; Heidaritabar *et al*. 2014).

At the functional level with model ADE, epistasis was abundant and 49% of the variation in the total genetic value was generated by functional epistatic effects and only 19% by functional additive effects. However, most of the genetic variance at the statistical level was additive (62%) or due to dominance (33%), and only 5% was epistatic variance in generation 0, which is reasonably close to results for litter size in pigs (Vitezica *et al*. 2018). The broad-sense heritability was set to 0.4 for all genetic models, resulting in a narrow-sense heritability of ∼0.25 for model ADE. This heritability was considerably lower than the narrow-sense heritability of ∼0.40 for model A and ∼0.38 for model AD. Altogether, those parameters indicate that the simulated genetic architecture using model ADE could represent the genetic architecture of a trait in a livestock population.

### Accuracy of selection

In the first generation of selection, the accuracy of selection was always highest with genomic selection including own performance (GBLUP_OP) (Figure 2). This accuracy was ∼0.83 for model A and model AD, and ∼0.72 for model ADE. This lower accuracy is a result of the lower narrow-sense heritability for this genetic model. For all genetic models, the accuracy of the pedigree selection scenario with own performance (PBLUP_OP) in generation 1 was ∼0.09 lower than with GBLUP_OP, the accuracy of genomic selection without own performance (GBLUP_NoOP) was ∼0.13 lower than with GBLUP_OP, and the accuracy of MASS was ∼0.21 lower than with GBLUP_OP. As expected, the accuracy of MASS was equal to the square root of the narrow-sense heritability. Over the generations, the accuracy of selection decreased for all scenarios. The decrease was largest in the first generations as a result of the Bulmer effect (Bulmer 1971). Thereafter, the decrease was slightly larger for the genomic selection scenarios (GBLUP_OP and GBLUP_NoOP) compared to PBLUP_OP and MASS. As a result, differences in accuracy between the scenarios after 50 generations of selection were smaller than in the first generation. The accuracy decreased fastest under genetic model ADE, especially for the genomic selection scenarios. Under this genetic model, the accuracies of PBLUP_OP, MASS and GBLUP_OP were similar after 50 generations of selection.

**FIGURE 2.**
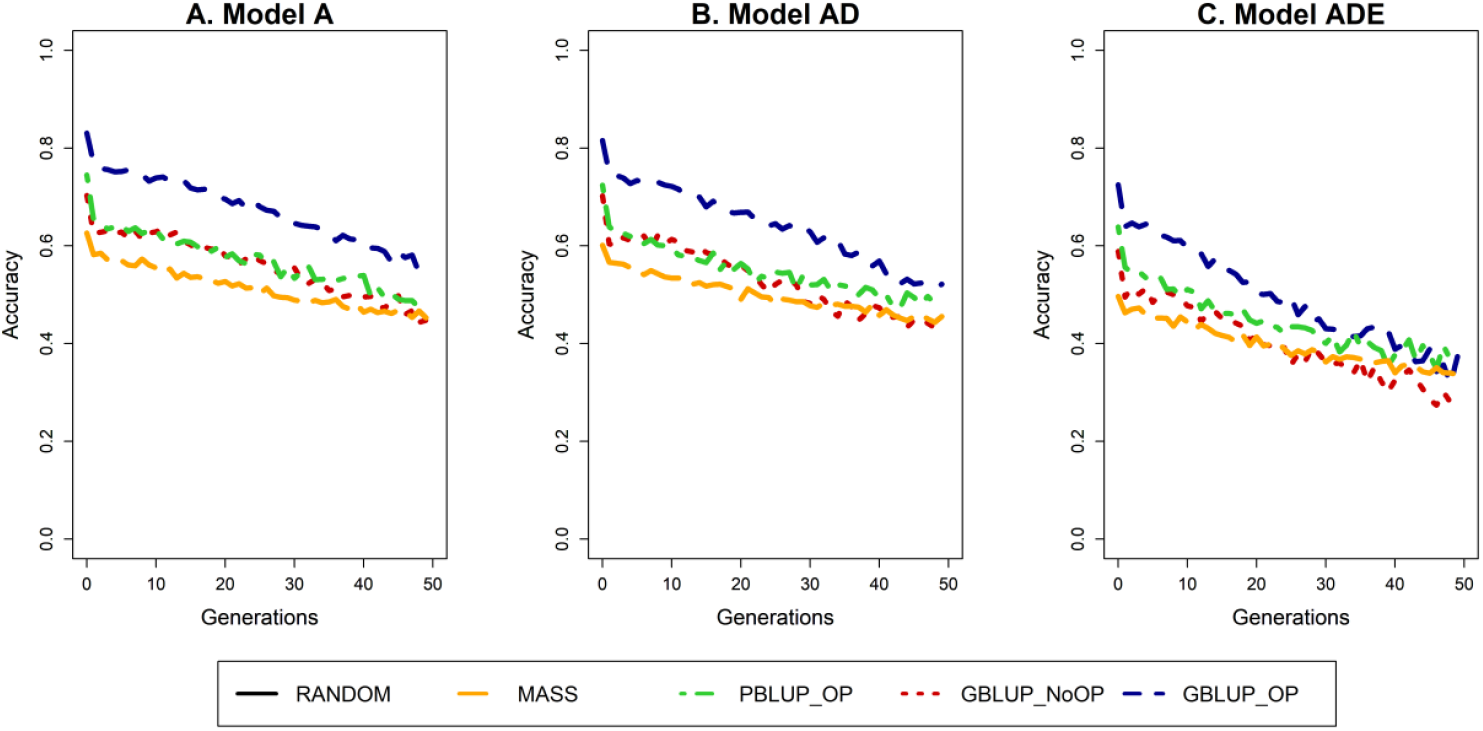
Trend in accuracy of selection for four selection methods and three genetic models. The four selection methods were: MASS selection, PBLUP selection with own performance (PBLUP_OP), GBLUP selection without own performance (GBLUP_NoOP) or with own performance (GBLUP_OP). The three genetic models were a model with only additive effects (A), with additive and dominance effects (AD), or with additive, dominance and epistatic effects (ADE). Results are shown as averages of 20 replicates.

### Genetic gain

Over generations, the average phenotypic value in the population was constant for the RANDOM scenario and increased with selection (Figure 3). The rates of genetic gain in the first generations resembled results for the accuracy, with highest values for GBLUP_OP, followed by PBLUP_OP, GBLUP_NoOP and finally MASS, and smaller values when non-additive effects were present. The rate of genetic gain decreased over generations, but considerably less for MASS than for the other selection methods. Therefore, after 50 generations of selection, MASS outperformed PBLUP_OP and GBLUP_NoOP in terms of accumulated genetic gain under all genetic models, and also outperformed GBLUP_OP under model ADE.

**FIGURE 3.**
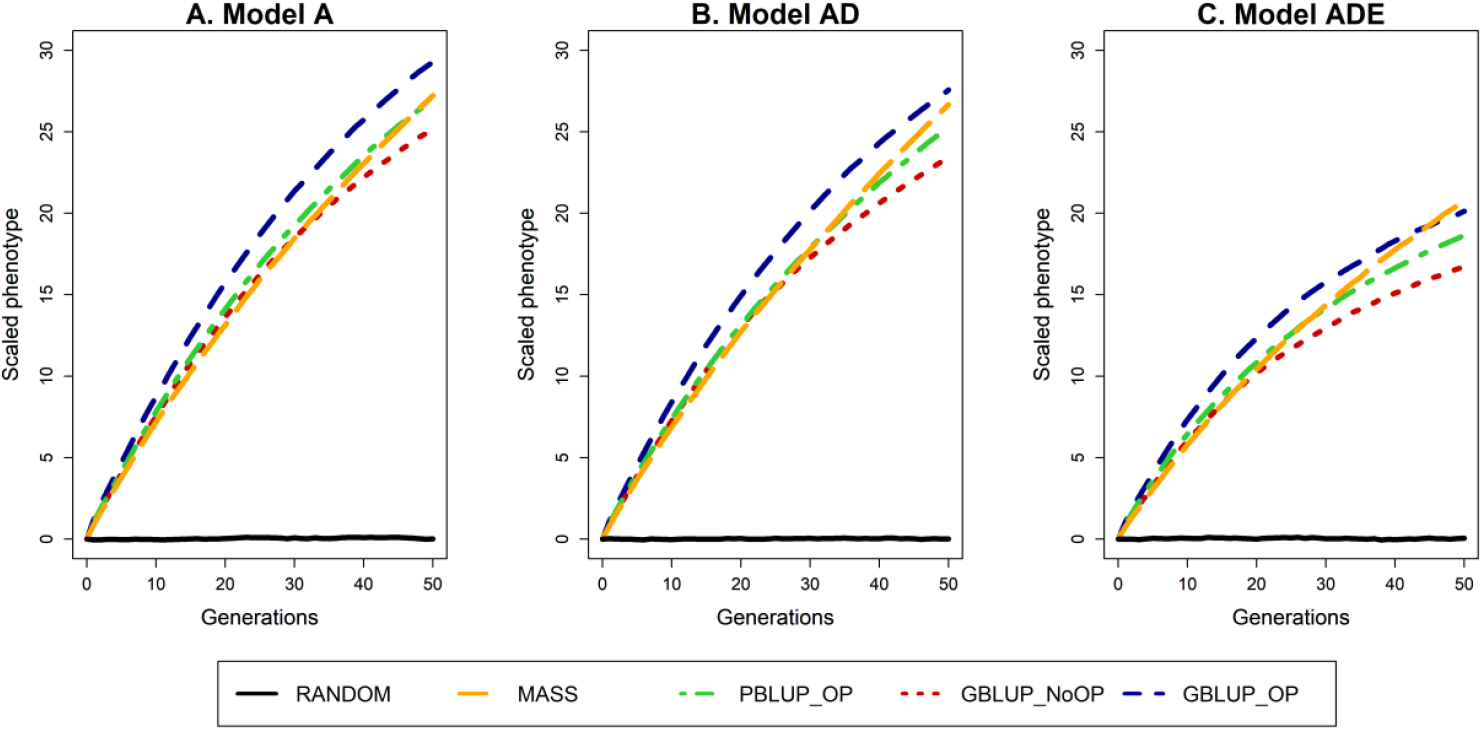
Phenotypic trend for the five selection methods and three genetic models. The phenotypic trend is scaled by the additive genetic standard deviation in the generation before selection in order to make the results comparable across the genetic models. The five selection methods were: RANDOM selection, MASS selection, PBLUP selection with own performance (PBLUP_OP), GBLUP selection without own performance (GBLUP_NoOP) or with own performance (GBLUP_OP). The three genetic models were a model with only additive effects (A), with additive and dominance effects (AD), or with additive, dominance and epistatic effects (ADE). Results are shown as averages of 20 replicates.

### Additive genetic and genic variance

The additive genetic and genic variance were approximately constant for the RANDOM scenario and decreased with selection (Figure 4). As expected, the largest drop in additive genetic variance was observed in the first generations of selection as a result of the Bulmer effect, as also observed for the accuracy of selection. In the first three generations of selection, genetic variance reduced by more than 20%. The total drop in genetic variance over 50 generations of selection was more or less similar for GBLUP_OP and GBLUP_NoOP where less than 20% of the initial genetic variance was maintained under genetic models A and AD. Under model ADE, more genetic variance was maintained (∼24%) after 50 generations of selection. Only slightly more genetic variance (∼25%) was maintained with PBLUP_OP, for which the loss in genetic variance was reasonably similar across the three genetic models. With MASS, the loss in genetic variance was considerably less, and ∼40% of the variance was maintained after 50 generations of selection.

**FIGURE 4.**
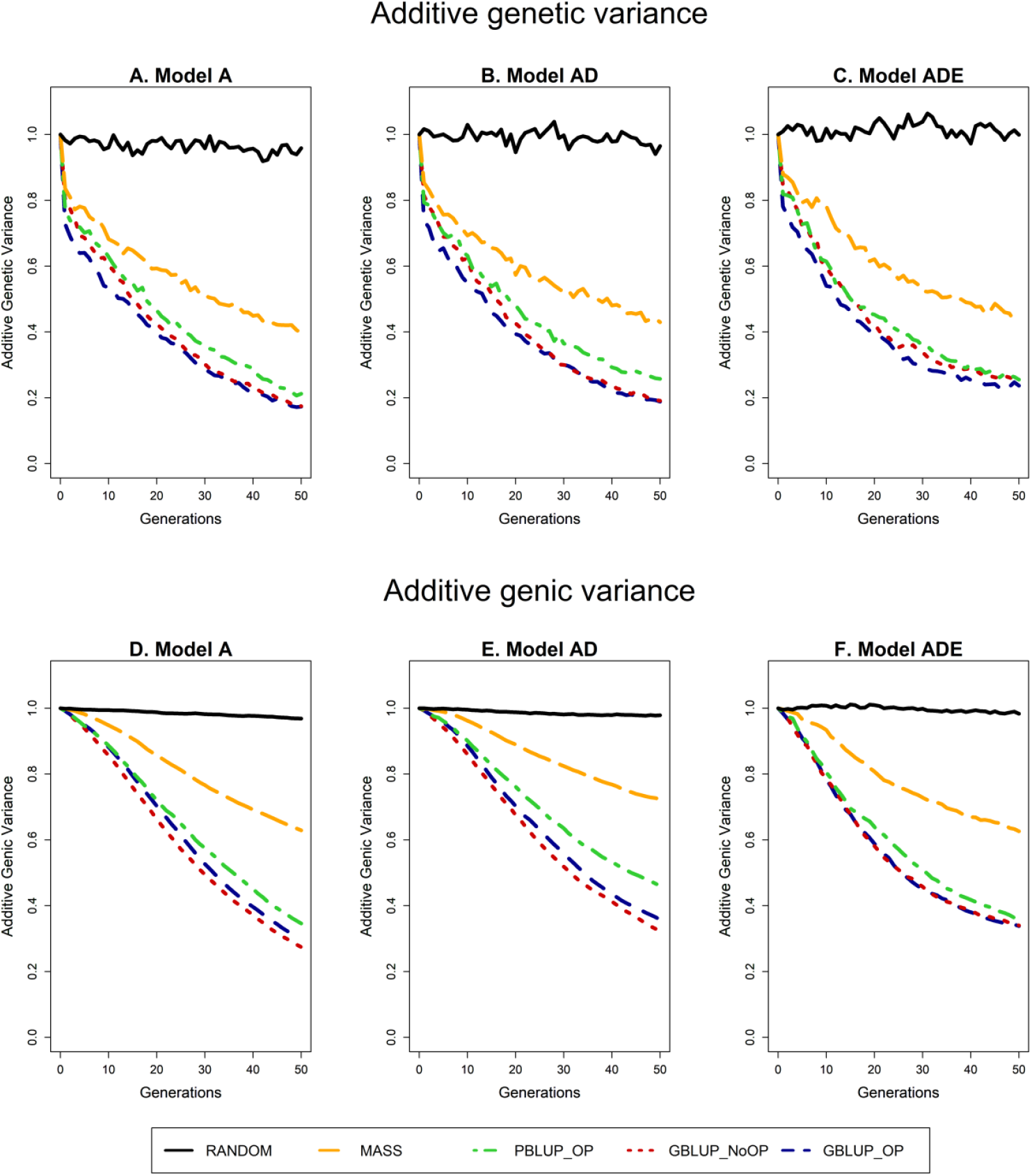
Trend in additive genetic (A, B, C) and additive genic (D, E, F) variance for the five selection methods and three genetic models. The trend is scaled by the additive genetic or additive genic variance in the generation before selection in order to make the results comparable across the genetic models. The five selection methods were: RANDOM selection, MASS selection, PBLUP selection with own performance (PBLUP_OP), GBLUP selection without own performance (GBLUP_NoOP) or with own performance (GBLUP_OP). The three genetic models were a model with only additive effects (A), with additive and dominance effects (AD), or with additive, dominance and epistatic effects (ADE). Results are shown as averages of 20 replicates.

The additive genic variance is not affected by transient effects such as the Bulmer effect (Bulmer 1971). Therefore, the loss in genic variance was smaller than the loss in genetic variance, especially in the first generations. Except for this difference in the first generations, the trends in additive genic and genetic variance were very similar.

### Number of segregating causal loci

The number of segregating causal loci decreased as a result of selection (Figure 5). For PBLUP_OP, the number of loci decreased fastest with a reduction of almost 50% over 50 generations of selection. For GBLUP_OP and GBLUP_NoOP the decrease was slightly smaller; 42% for GBLUP_OP and 40% for GBLUP_NoOP. For MASS, the decrease was substantially smaller and the number of loci decreased by only 20%. The loss in segregating loci was slightly smaller when non-additive effects were present. Interestingly, the number of segregating loci in generation 50 was lower for PBLUP_OP than for GBLUP_OP and GBLUP_NoOP, while the additive genic variance was slightly larger for PBLUP_OP.

**FIGURE 5.**
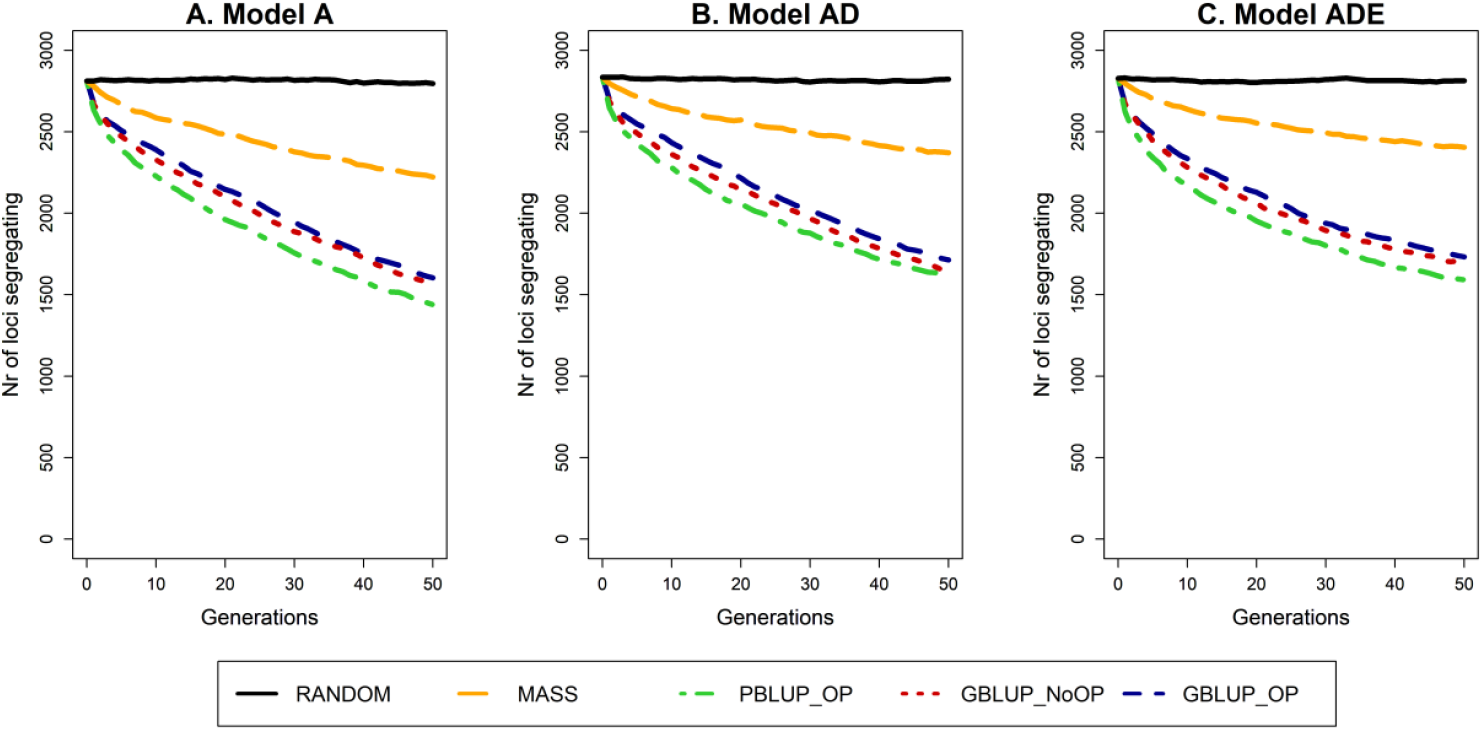
Trend in number of segregating causal loci for the five selection methods and three genetic models. The five selection methods were: RANDOM selection, MASS selection, PBLUP selection with own performance (PBLUP_OP), GBLUP selection without own performance (GBLUP_NoOP) or with own performance (GBLUP_OP). The three genetic models were a model with only additive effects (A), with additive and dominance effects (AD), or with additive, dominance and epistatic effects (ADE). Results are shown as averages of 20 replicates.

### Average minor allele frequency at segregating causal loci

The additive genic variance depends on the number of segregating causal loci as well as their MAF. In the first generations of selection, the average MAF of segregating loci increased, especially for PBLUP_OP (Figure 6). Thereafter, the average MAF decreased and after 50 generations of selection, it was below its initial value, with the smallest values for the GBLUP scenarios. Only under genetic model ADE, the average MAF of PBLUP_OP and MASS after 50 generations of selection were slightly above the average MAF before selection. The impact of MASS on the average MAF of segregating loci was very limited. The higher average MAF of PBLUP_OP can explain the higher additive genic variance for PBLUP_OP than for GBLUP_OP and GBLUP_NoOP, even though the number of segregating loci was smaller (Figure 4 vs 5).

**FIGURE 6.**
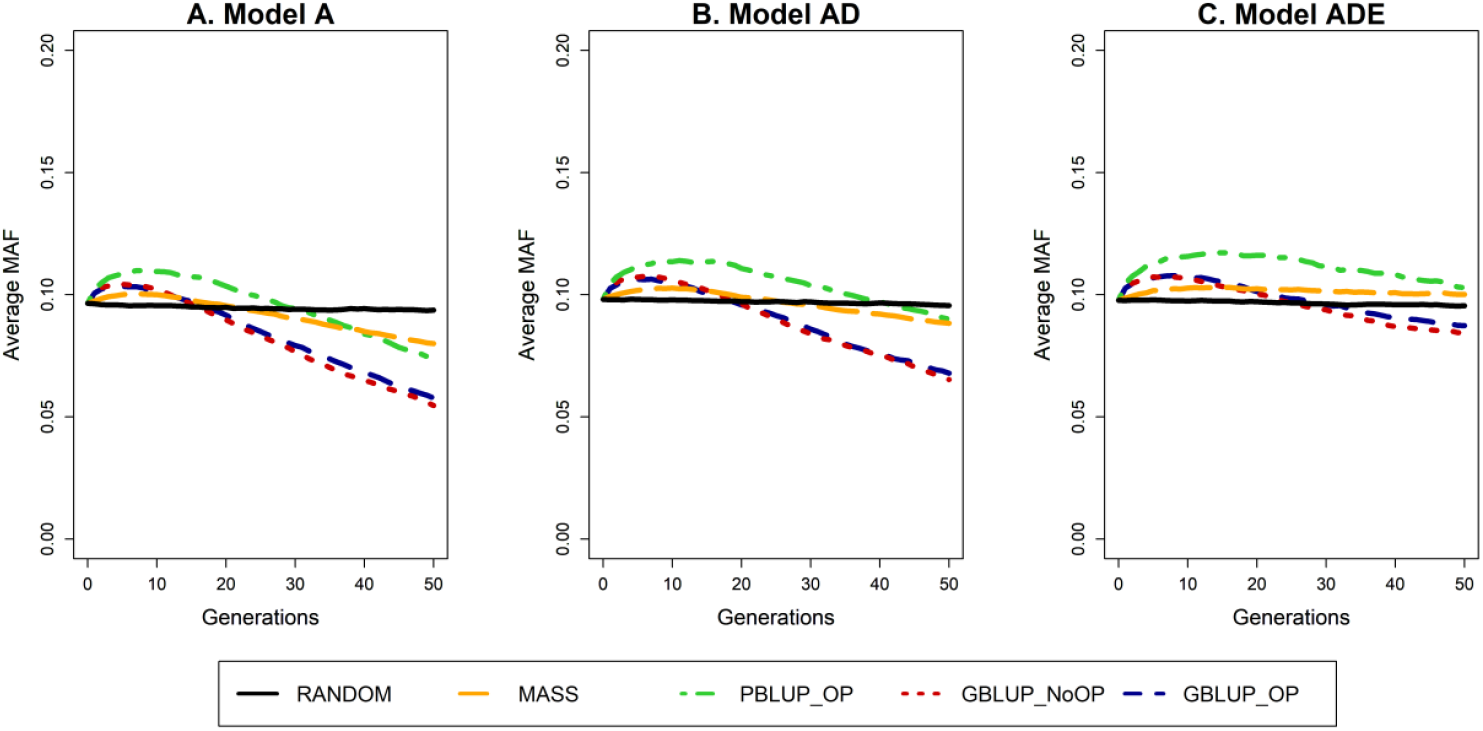
Trend in average minor allele frequency (MAF) of segregating causal loci for the five selection methods and three genetic models. The five selection methods were: RANDOM selection, MASS selection, PBLUP selection with own performance (PBLUP_OP), GBLUP selection without own performance (GBLUP_NoOP) or with own performance (GBLUP_OP). The three genetic models were a model with only additive effects (A), with additive and dominance effects (AD), or with additive, dominance and epistatic effects (ADE). Results are shown as averages of 20 replicates.

### Accumulated heterozygosity

In a random mating population, the accumulated heterozygosity depends on the number of segregating causal loci (Figure 5), their average MAF (Figure 6) and the variation in MAF among loci (Supplemental file 2, Figure S2.3 and Supplemental file 3). As expected, selection resulted in a decrease in the accumulated heterozygosity (Figure 7). The reduction in accumulated heterozygosity was similar for GBLUP_OP and GBLUP_NoOP, slightly less for PBLUP_OP and considerably less for MASS. Moreover, the accumulated heterozygosity decreased slower when non-additive effects were present. Thus, the decrease in heterozygosity was lower for pedigree than for genomic selection, and depended on the genetic model.

**FIGURE 7.**
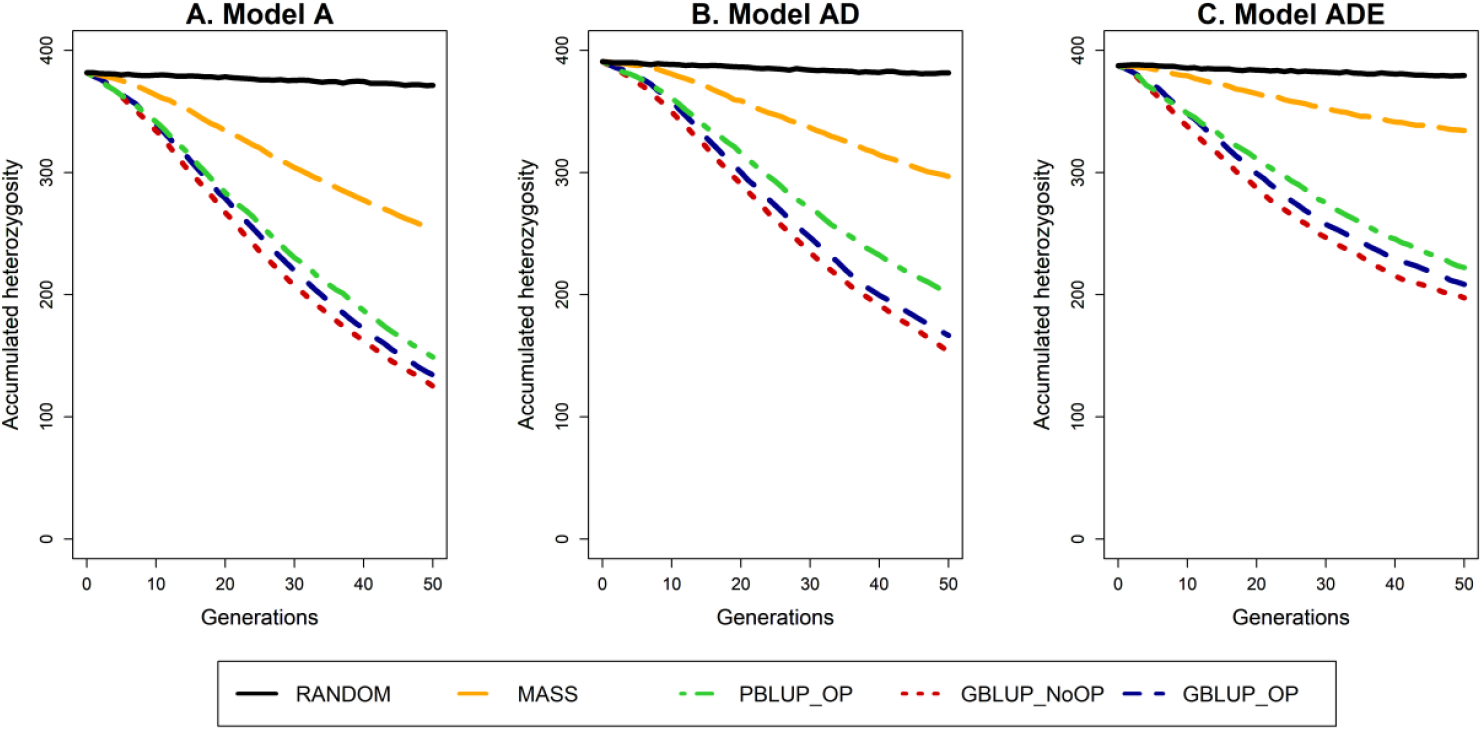
Trend in accumulated heterozygosity across segregating causal loci for the five selection methods and three genetic models. The five selection methods were: RANDOM selection, MASS selection, PBLUP selection with own performance (PBLUP_OP), GBLUP selection without own performance (GBLUP_NoOP) or with own performance (GBLUP_OP). The three genetic models were a model with only additive effects (A), with additive and dominance effects (AD), or with additive, dominance and epistatic effects (ADE). Results are shown as averages of 20 replicates.

### Change in genetic architecture

Over generations, the subset of causal loci underlying the trait (Figure 8), the allele frequencies of the causal loci (Figure 9), and the statistical additive effects of the causal loci (Figure 10) changed. The change in the subset of loci was measured by the Jaccard index. Especially in the first generation, the subset of loci changed considerably, because every generation had approximately 600 new mutations, most of which were lost immediately. As a result, two consecutive generations already differ in almost 1200 loci. The subset of loci affecting the trait changed considerably with drift (RANDOM), but the change was amplified by selection. After 50 generations, the average Jaccard index was ∼0.27 for RANDOM, ∼0.21 for MASS and between 0.10 and 0.15 for PBLUP_OP, GBLUP_NoOP, and GBLUP_OP. The Jaccard index was slightly higher with non-additive effects. Those results indicate that the subset of loci affecting the trait constantly changes over generations due to new mutations and drift, and that the change is amplified by selection.

**FIGURE 8.**
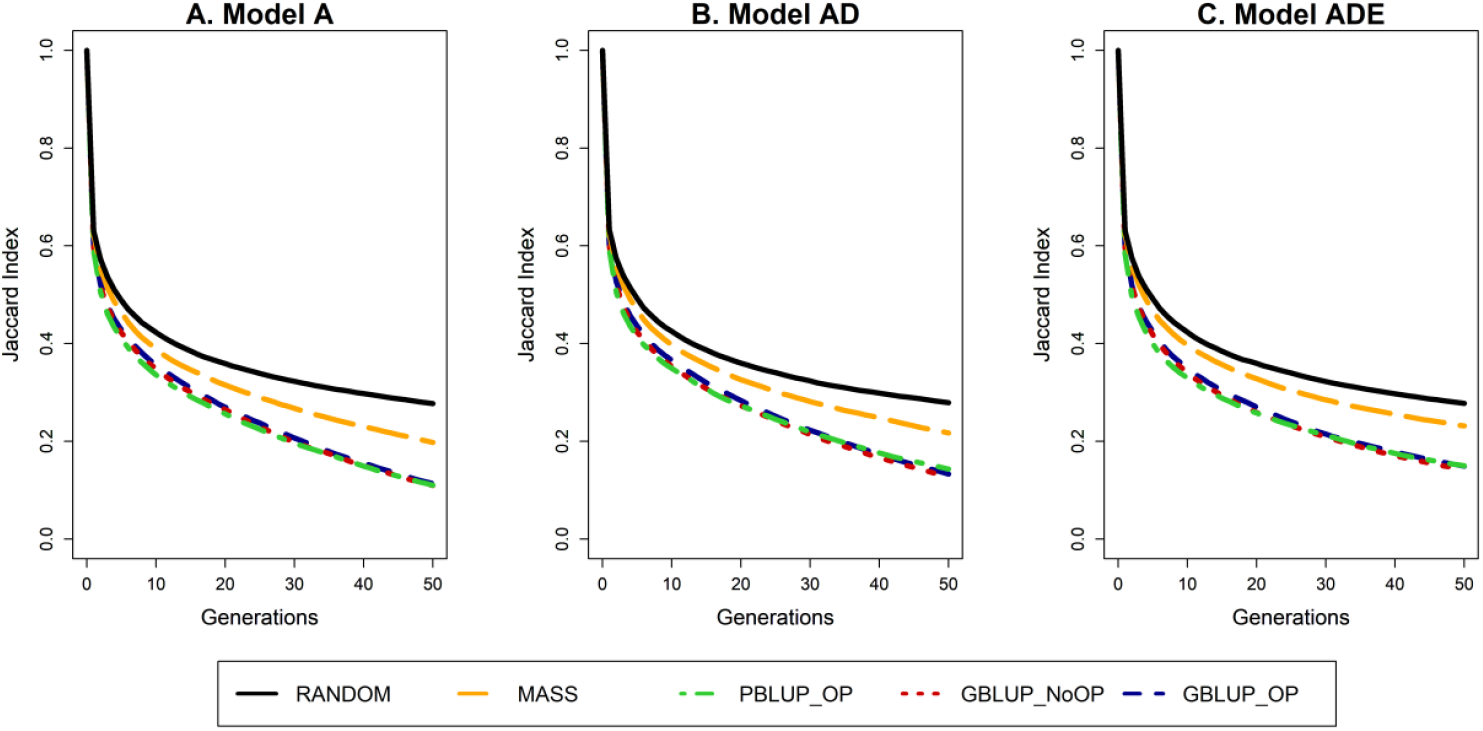
Change in the subset of segregating causal loci for the five selection methods and three genetic models. The change in the subset is described by the Jaccard index. The five selection methods were: RANDOM selection, MASS selection, PBLUP selection with own performance (PBLUP_OP), GBLUP selection without own performance (GBLUP_NoOP) or with own performance (GBLUP_OP). The three genetic models were a model with only additive effects (A), with additive and dominance effects (AD), or with additive, dominance and epistatic effects (ADE). Results are shown as averages of 20 replicates.

**FIGURE 9.**
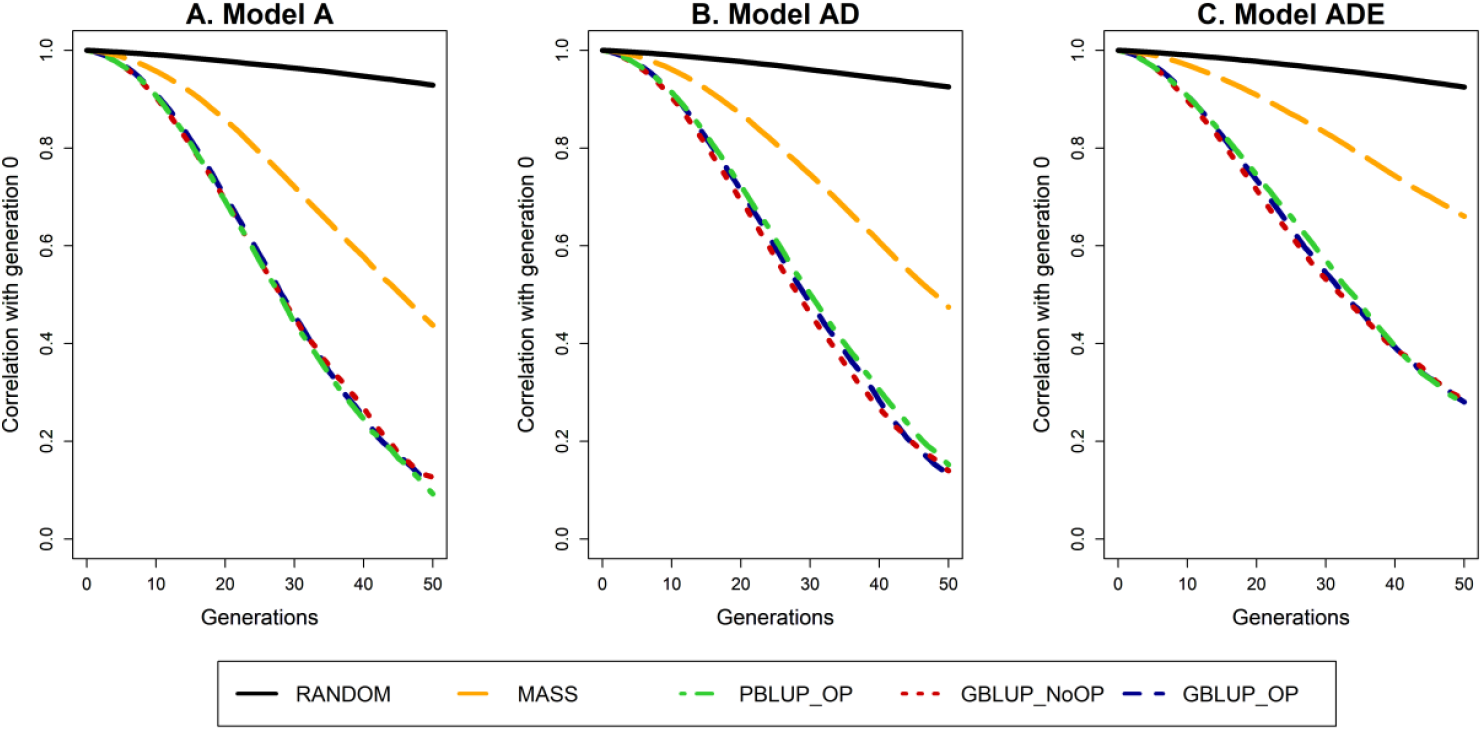
Change in the allele frequencies of segregating causal loci for the five selection methods and three genetic models. The change in allele frequencies is represented by the correlation in allele frequencies between the generation of interest and the generation before selection (generation 0). The five selection methods were: RANDOM selection, MASS selection, PBLUP selection with own performance (PBLUP_OP), GBLUP selection without own performance (GBLUP_NoOP) or with own performance (GBLUP_OP). The three genetic models were a model with only additive effects (A), with additive and dominance effects (AD), or with additive, dominance and epistatic effects (ADE). Results are shown as averages of 20 replicates.

**FIGURE 10.**
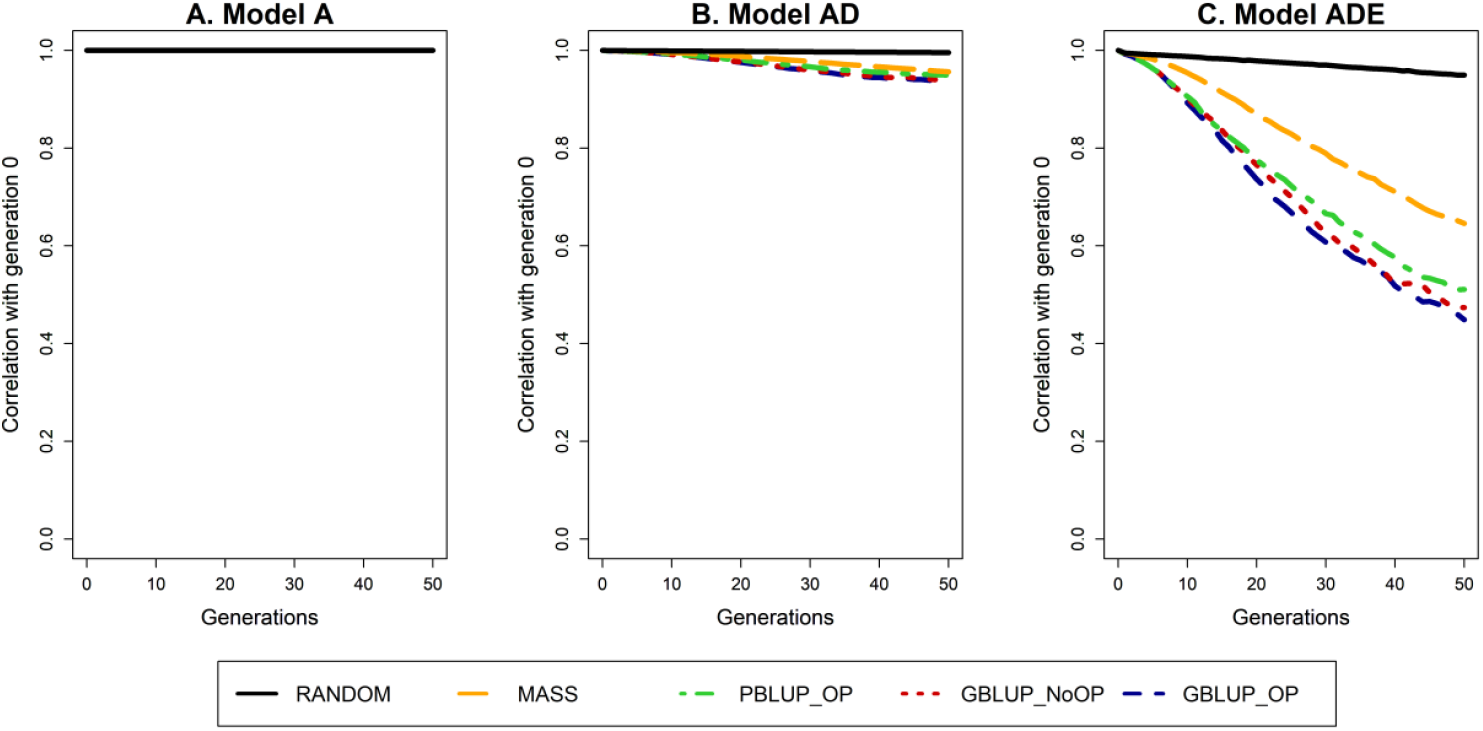
Change in the statistical additive effects of segregating causal loci for the five selection methods and three genetic models. The change in statistical additive effects is represented by the correlation in the effects between the generation of interest and the generation before selection (generation 0). The five selection methods were: RANDOM selection, MASS selection, PBLUP selection with own performance (PBLUP_OP), GBLUP selection without own performance (GBLUP_NoOP) or with own performance (GBLUP_OP). The three genetic models were a model with only additive effects (A), with additive and dominance effects (AD), or with additive, dominance and epistatic effects (ADE). Results are shown as averages of 20 replicates.

Selection strongly amplified the change in allele frequencies of loci compared to drift (Figure 9; Supplemental file 4). Due to drift alone, the correlation between the allele frequencies of loci segregating in both generation 0 and generation 50 was ∼0.93 (RANDOM). The change in allele frequencies as a result of selection was largest under model A, with a correlation between the allele frequencies of generation 0 and 50 of only ∼0.10 for GBLUP_OP, GBLUP_NoOP and PBLUP_OP, and of 0.44 for MASS. Those correlations were slightly higher under model AD. When also epistatic effects were present, the change in allele frequencies was much smaller, and the correlation was ∼0.28 after 50 generations of GBLUP_OP, GBLUP_NoOP and PBLUP_OP, and 0.66 for MASS.

As a result of the change in allele frequency, statistical additive effects of the loci changed when non-additive effects were present (Figure 10; Supplemental file 5). The changes were quite limited when only additive and dominance effects were present, with a correlation of ∼0.94 between the statistical additive effects of generation 0 and 50 for all selection methods. When epistatic effects were also present, this correlation was much lower. After 50 generations, the average correlation was 0.95 for RANDOM, 0.65 for MASS, 0.51 for PBLUP_OP, 0.47 for GBLUP_NoOP, and 0.45 for GBLUP_OP. Within 10 generations of GBLUP_OP, GBLUP_NoOP or PBLUP_OP, the correlation had already dropped to ∼0.90.

## DISCUSSION

We investigated the long-term effects of genomic selection on the rate of genetic gain, additive genetic variance and genetic architecture of traits. Over 50 generations of genomic selection (GBLUP), the accuracy of selection, the rate of genetic gain, the amount of additive genetic and genic variation, and the number of segregating causal loci decreased. The same trends were also observed for phenotypic (MASS) and pedigree (PBLUP) selection, but the decrease was considerably smaller for MASS and slightly smaller for PBLUP. The main results of our study are assembled in Table 2, which also mentions the most likely mechanism underlying the results that will be further explained in this discussion.

**Table 2.**
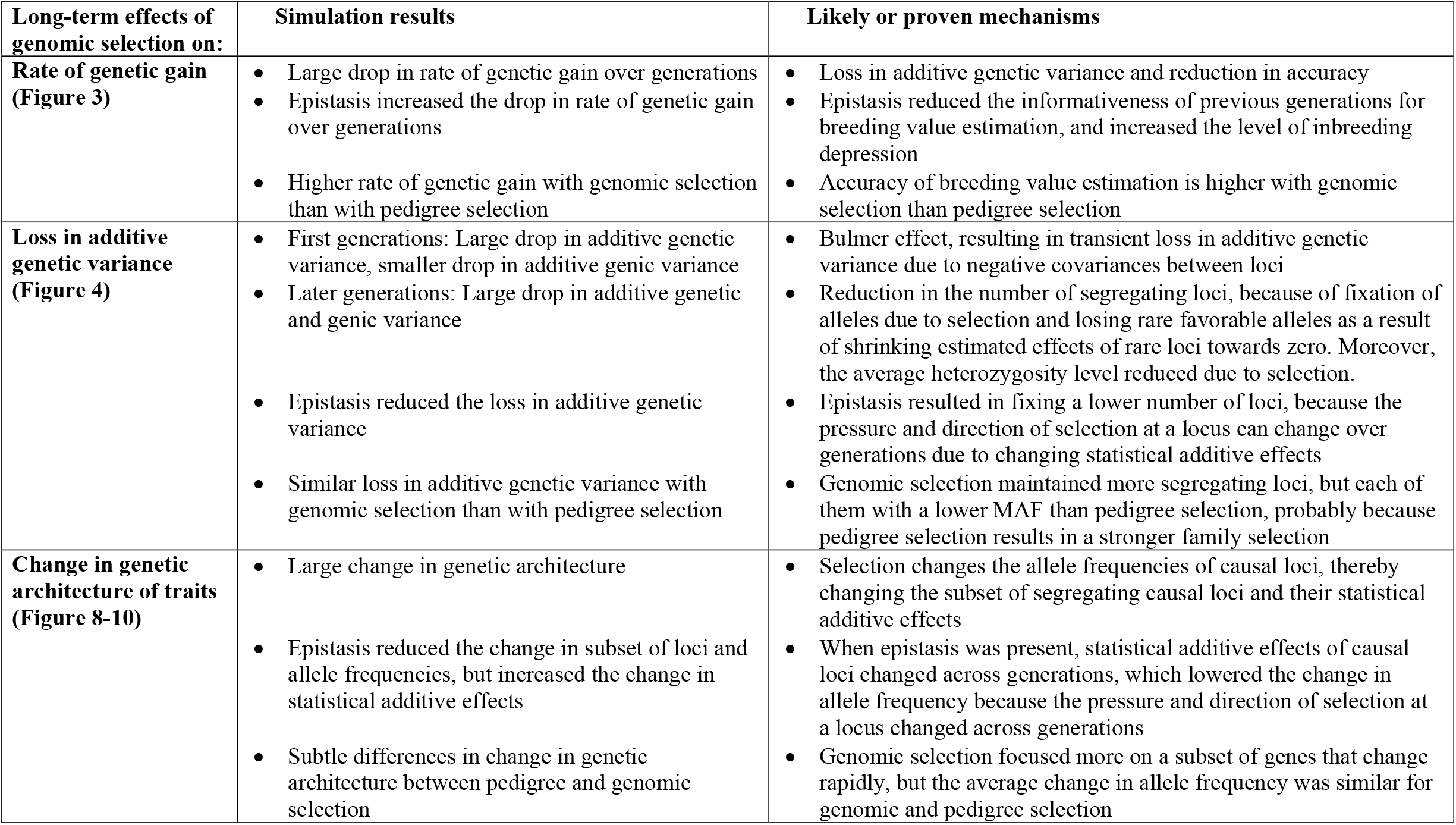
Summary table of long-term effects of genomic selection

### Genetic gain

MASS yielded the lowest initial rate of genetic gain, but the highest rate of genetic gain after 50 generations (Figure 3). Therefore, MASS outperformed most of the other selection methods in terms of cumulative genetic gain over 50 generations of selection. Those results are in agreement with previous research based on the infinitesimal model, which showed that MASS can outperform PBLUP for long-term gain, even though short-term gain was highest for PBLUP (Verrier *et al*. 1993; Wei *et al*. 1996). This was mainly because PBLUP lost more genetic variation (Figure 4) and segregating loci (Figure 5) than MASS.

The relative benefit of MASS for cumulative genetic gain over 50 generations was larger with dominance. The other selection methods that each had a higher initial accuracy than MASS resulted in selecting more related individuals; the pedigree inbreeding coefficients after 50 generations was ∼2.6 times larger with PBLUP and ∼1.8 times larger with GBLUP than with MASS (Supplemental file 6, Table S6.1). These larger inbreeding coefficients with PBLUP and GBLUP suggests that those selection methods suffered more from inbreeding depression than MASS in the presence of dominance. It has been shown previously that with high rates of inbreeding depression, the long-term genetic gain is higher with MASS than with PBLUP (Quinton *et al*. 1992).

The benefit of MASS in cumulative genetic gain was even larger when also epistasis was present. This is a result of the increasingly smaller difference in accuracy between selection methods over generations when epistasis was present than when epistasis was absent, because the accuracy of GBLUP and PBLUP dropped faster over generations when epistasis was present (as will be further explained later). The smaller difference in accuracy between selection methods together with the much higher additive genetic variance in the last generations for MASS (Figure 4) resulted in a much higher rate of genetic gain for MASS in those generations compared to the other selection methods.

The cumulative genetic gain was always lowest for GBLUP without own performance records (GBLUP_NoOP). This selection method, however, allows for a considerable reduction in the generation interval in some species, because selection can take place at a younger age before own phenotypic information or progeny information is available. This is most pronounced in dairy cattle, where the generation interval can be halved (Schaeffer 2006; García-Ruiz *et al*. 2016). In this study, a potential difference in generation interval was not taken into account because our focus is on the genetic mechanisms, not on applied breeding programs.

### Genetic variance

All selection methods resulted in a significant loss in genetic variance (Figure 4). Part of this loss was transient and a result of the Bulmer effect (Bulmer 1971). The difference between the genetic and genic variance was, however, reasonably small (Supplemental file 2, Figure S2.4). This small difference indicates that the largest part of the loss in genetic variance was a result of allele frequency changes, and thus permanent.

Genic variance was lost across all 50 generations of selection and here we investigate the trends in the different components of the genic variance; number of segregating causal loci (*n*), average heterozygosity 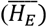, average square of the statistical additive effects 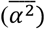 and covariance between heterozygosity and α^2^ (*Cov(H*_*E*_, α^2^); Supplemental file 3). In the first generations, genic variance was lost due to a considerable drop in the number of segregating loci, which was slightly counteracted by an increase in 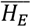 at those loci (Supplemental file 6, Table S6.2). Especially with PBLUP, 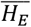 increased in the first generations of selection. After generation 10, 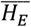 decreased and was lower after 50 generations than before selection for most scenarios. In generation 50, 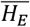 was higher for both MASS and PBLUP than for GBLUP. Moreover, loci with a larger statistical additive effect were more likely to become fixed, which slightly reduced 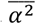, especially when epistasis was present. The covariance between *H*_*E*_ and α^2^ was in general close to zero and contributed only little to the genic variance. Altogether, those results show that after 50 generations of selection, the drop in genic variance could for the largest part be explained by a reduction in the number of segregating loci and in 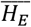 at those loci, and for a smaller part by a reduction in 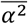.

The total loss in genic variance was lowest for MASS, which maintained a much higher number of segregating loci than PBLUP or GBLUP (Figure 5, Supplemental file 6, Table S6.2). This is probably because MASS better exploits rare favorable alleles. In GBLUP and maybe implicitly and for a smaller extent also in PBLUP, the effects of rare alleles are heavily regressed towards zero because they contribute little to the genetic variance (Goddard 2009; Gianola 2013). This enlarges the risk of keeping those alleles at low frequency or of losing them (Jannink 2010; Liu *et al*. 2015; De Beukelaer *et al*. 2017), even though they have the potential to greatly contribute to the genetic variance and future genetic gain. The number of rare alleles (MAF < 0.01) that was lost or remained rare during 50 generations of selection was indeed lower with MASS than with GBLUP and PBLUP (Supplemental file 6, Table S6.3). Therefore, the shrinkage of effects of rare alleles in PBLUP and GBLUP is the most likely explanation for losing more rare favorable alleles than with MASS.

The total loss in genic variance was comparable for GBLUP and PBLUP. The number of segregating loci, however, decreased faster with PBLUP, while the average MAF of segregating loci and thereby the heterozygosity level was higher with PBLUP (Figure 6; Supplemental file 6, Table S6.2). This might be a result of a stronger family selection with PBLUP, which agrees with the higher pedigree inbreeding level observed with PBLUP (Supplemental file 6, Table S6.1).

We also investigated the impact of this difference beyond the 50 generations we simulated by estimating for each scenario the theoretical maximum genetic gain that can still be achieved from generation 50 onwards. This maximum genetic gain would be achieved when all loci would become fixed for the favourable allele, using the statistical additive effects of generation 50 (Supplemental file 6, Table S6.4). The maximum genetic gain was on average 7.6% and 6.1% higher for GBLUP with or without own performance compared to PBLUP with own performance. This suggests that GBLUP is more sustainable for maintaining future genetic gain than PBLUP.

The loss in genic variance was slightly smaller with non-additive effects. With non-additive effects, the statistical additive effects depend on the allele frequencies (Fisher 1930; Mackay 2014). Some positive statistical additive effects even changed into negative effects over generations when epistasis was present (Supplemental file 5), which also changed the direction of selection. Those changes limited the number of loci that became fixed in the population, because selection was not constantly focussing on the same alleles or loci across generations. This resulted in a higher number of segregating loci (Figure 5) and a higher level of accumulated heterozygosity (Figure 7) after 50 generations of selection when non-additive effects were present (Supplemental file 6, Table S6.2).

### Genetic architecture

Our initial plan was to quantify the change in genetic architecture of traits by the genetic correlation between generations. However, this turned out to be very complex. The genetic correlation is defined as the correlation between the additive genetic values (i.e., true breeding values) for two traits of the same individual (Bohren *et al*. 1966; Falconer and Mackay 1996). For a genetic correlation between generation 1 and 10, for example, trait 1 reflects the true breeding value in generation 1 and trait 2 the true breeding value in generation 10. Due to the change in allele frequencies over generations, the correlation between generation 1 and 10 would be different when breeding values were estimated for a set of individuals from generation 1, generation 10, or both generation 1 and 10 (Duenk *et al*. 2020). This means that the genetic correlation between generations depends on the subset of individuals used for estimating the genetic correlation, and is not influenced by the change in allele frequencies and subset of loci underlying the trait across generations. Therefore, we decided to quantify the change in genetic architecture across generations using three measures based on the underlying mechanisms; the change in subset of loci, allele frequencies and statistical additive effects.

The change in genetic architecture was strongly enhanced by selection, because selection resulted in a faster changes in the subset of causal loci, in allele frequencies and in statistical additive effects (Figures 8, 9 and 10). Contrary to our expectation and earlier results (Heidaritabar *et al*. 2014; Liu *et al*. 2014), the change in genetic architecture was about similar for GBLUP and PBLUP. This indicates that even though GBLUP focusses more on a subset of the genome that changes rapidly in allele frequencies while PBLUP spreads the selection pressure more evenly across the genome, the average change in allele frequencies was about equal. This was confirmed by the larger variance in the change in allele frequency at loci for GBLUP than PBLUP (Supplemental file 6, Table S6.5).

After five generations of GBLUP and PBLUP, the Jaccard index had already dropped to 0.42, the correlation between allele frequencies was 0.97, and the correlation between statistical additive effects 0.96 under model ADE. These results were very similar when they were calculated with generation 5 as reference instead of generation 0 (Supplemental file 2, Figures S2.5 – S2.7), indicating that the rate of change in genetic architecture was not higher at the start of selection than in later generations. We hypothesized that those changes could result in changes in true breeding values across generations. Therefore, we estimated the correlation in true breeding values of individuals from generation 50 for performance in generation 50 and generations 47 to 49 that were included in the reference population. The correlation between true breeding values was always >0.99 when only additive or additive and dominance effects were present, but substantially smaller than 1 (∼0.95 with generation 49, ∼0.91 with generation 48, and ∼0.87 with generation 47) for the PBLUP and GBLUP scenarios with epistasis (Table 3). This indicates that even though the correlation in statistical additive effects was very high between neighboring generations (>0.99, Figure 10), the correlation of true breeding values between generations decreased rapidly because statistical additive effects changed more rapidly for loci with high MAF or large effect (Supplemental file 5). This phenomenon drastically decreased the informativeness of previous generations for breeding value prediction. So, recent generations of reference populations for genomic prediction are more useful, not only because they are closer related to the selection candidates (Clark *et al*. 2012; Pszczola *et al*. 2012; Wientjes *et al*. 2013), but also because their genetic architecture is more similar to that in the selection candidates.

**TABLE 3.** Correlation of true breeding values (TBV) of individuals from generation 50 between performance in generation 50 and each of the three previous generations.

|  | **Correlation in TBV of generation 50 with** |  |  |
| --- | --- | --- | --- |
|  | **Generation 49** | **Generation 48** | **Generation 47** |
| *Model A* |  |  |  |
| **RANDOM** | 1.00 | 1.00 | 1.00 |
| **MASS** | 1.00 | 1.00 | 1.00 |
| **PBLUP** | 1.00 | 1.00 | 1.00 |
| **GBLUP_NoOP** | 1.00 | 1.00 | 1.00 |
| **GBLUP_OP** | 1.00 | 1.00 | 1.00 |
| *Model AD* |  |  |  |
| **RANDOM** | 1.00 | 1.00 | 1.00 |
| **MASS** | 1.00 | 1.00 | 1.00 |
| **PBLUP** | 1.00 | 0.99 | 0.99 |
| **GBLUP_NoOP** | 1.00 | 0.99 | 0.99 |
| **GBLUP_OP** | 1.00 | 0.99 | 0.99 |
| *Model ADE* |  |  |  |
| **RANDOM** | 0.99 | 0.99 | 0.99 |
| **MASS** | 0.98 | 0.98 | 0.97 |
| **PBLUP** | 0.95 | 0.92 | 0.90 |
| **GBLUP_NoOP** | 0.96 | 0.91 | 0.86 |
| **GBLUP_OP** | 0.95 | 0.91 | 0.87 |

### Non-additive effects

The contribution of epistasis to the variation in quantitative traits is a highly debated topic. The results of our study show that even though almost 50% of the variation in the total genetic value was generated by functional epistatic effects, the epistatic genetic variance explained only 5% of the genetic variance. This means that most of the epistatic effects were captured in the statistical additive effects, which was also expected based on the U-shaped allele frequency distribution of the loci (Hill *et al*. 2008; Mäki-Tanila and Hill 2014). So the fact that populations only show a limited amount of statistical epistatic variance does not prove that the amount of functional epistasis in the population is limited (Cheverud and Routman 1995; Huang and Mackay 2016).

Depending on allele frequencies, a large part of the functional dominance and epistatic effects can be converted into additive variance (Barton and Turelli 2004; Carlborg *et al*. 2006; Le Rouzic and Carlborg 2008; Hill 2017). For the model with additive and dominance effects, 96% of the additive genetic variance was a result of functional additive effects before selection, and roughly all genetic variance after 50 generations of selection (Supplemental file 2, Figure S2.8). When epistatic effects were also present, only 34% of the additive genetic variance was a result of functional additive effects before selection and roughly 50% after selection. This shows that the part of the functional dominance and epistatic effects captured by the statistical additive variance changes across generations due to allele frequency changes. The conversion of non-additive effects into statistical additive effects depends on the allele frequencies. When allele frequencies are closer to 0 or 1, a larger proportion of the non-additive effects is converted into statistical additive effects. As a result, a negative correlation between MAF and the absolute statistical additive effect of a locus existed in our simulations already before selection, even though functional effects were simulated independently of allele frequencies (Figure 11). A negative correlation between MAF and the effect size of loci is often observed in empirical studies (Manolio *et al*. 2009; Marouli *et al*. 2017; Zeng *et al*. 2018), and our results show that the existence of non-additive effects can contribute to explaining this finding.

**FIGURE 11.**
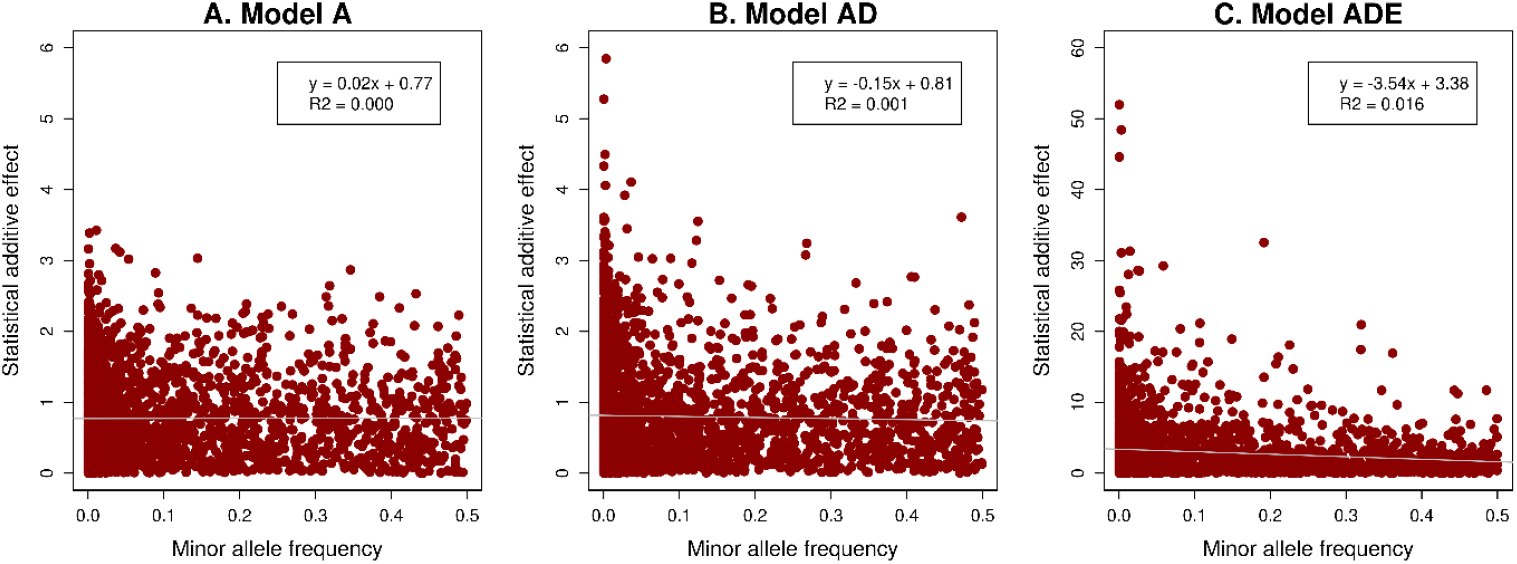
Correlation between the absolute statistical additive effect and minor allele frequency at causal loci for the three genetic models. The three genetic models were a model with only additive effects (A), with additive and dominance effects (AD), or with additive, dominance and epistatic effects (ADE).

Little is known about the structure and network of epistatic interactions. We only simulated pairwise interactions and mimicked the genetic interaction network described in yeast, with many loci with few interactions and few loci with many interactions. Similar interaction networks although studied in less detail are found in *C. Elegans* (Lehner *et al*. 2006), *Drosophila* (Huang *et al*. 2012) and mice (Tyler *et al*. 2017), and are also found between proteins (Tong *et al*. 2004). Therefore, Boone *et al*. (2007) and Mackay (2014) argue that it is likely that the interaction network between genetic loci is similar in other species such as livestock and human as well.

## Conclusion

An overview of the main results of this study is shown in Table 2. Our results show that GBLUP with own performance records resulted in the highest short-term genetic gain, while long-term gain was highest with MASS. This was mainly a result of a much higher loss in genetic variance and number of segregating loci with GBLUP. GBLUP without own performance records showed a slightly higher short-term gain than MASS, but considerably lower long-term gain. The genetic gain of PBLUP with own performance records was in between GBLUP with and without own performance records. PBLUP and GBLUP showed a similar loss in genetic variance, but the underlying mechanism was different; GBLUP maintained more loci, but with a lower MAF. The maximum genetic gain that could still be obtained after 50 generations of GBLUP selection was higher, which suggests that GBLUP better maintains long-term genetic gain than PBLUP. We have also shown that the change in genetic architecture of traits was strongly amplified by selection, with larger changes in the subset, allele frequencies and statistical additive effects of loci. However, in contrast to our hypothesis, the rate of change in genetic architecture was comparable for genomic and pedigree selection. Moreover, our results show that non-additive effects were relatively unimportant in the short-term, but they can substantially impact the accuracy and genetic gain of genomic selection when multiple generations are included in the reference population.

## ACKNOWLEDGMENTS

This research is supported by the Netherlands Organisation of Scientific Research (NWO; VENI grant 16774). The use of the HPC cluster has been made possible by CAT-AgroFood (Shared Research Facilities Wageningen UR).

